# Epigenetic Effects of Assisted Reproductive Technology in Human Offspring

**DOI:** 10.1101/816157

**Authors:** Wei Chen, Yong Peng, Xinyi Ma, Siming Kong, Shuangyan Tang, Yuan Wei, Yangyu Zhao, Wenxin Zhang, Yang Wang, Liying Yan, Jie Qiao

## Abstract

The births of more than 8 million infants have been enabled globally through assisted reproductive technologies (ARTs), including conventional in vitro fertilization (IVF) and intracytoplasmic sperm injection (ICSI) with either fresh embryo transfer (ET) or frozen embryo transfer (FET). However, the potential for elevated risks of ART-related disorders persists in adult life, and the underlying epigenetic mechanisms are largely uncharacterized. Here, we recruited 100 nuclear families and profiled the DNA methylomes, genome-wide histone modifications and transcriptomes to clarify the inherent extra risks attributable to specific ART procedures. We discovered that IVF-ET seemed to introduce less disturbance into the infant epigenome than IVF-FET or ICSI-ET did. Furthermore, we noted approximately half of the DNA methylomic changes in ART-conceived offspring could be explained by parental background biases. Through removal of the parental effect, we confirmed that ART *per se* would introduce minor DNA methylation changes locally. More importantly, we found that ART-induced epigenomic alterations were highly enriched in the processes which might contribute to increased incidence of preeclampsia during pregnancy and metabolic syndrome in offspring. Overall, our study provides an epigenetic basis for the potential long-term health risks in ART-conceived offspring that reinforces the need to review all methods of human ART.

## Introduction

Assisted reproductive technology (ART) has become routine in infertile treatment; indeed, more than eight million ART-conceived infants have been born worldwide^1^. However, conventional in vitro fertilization and fresh embryo transfer (IVF-ET) will introduce extraordinary changes in the environment where oocytes mature and the early embryo develops^2^. Moreover, intracytoplasmic sperm injection (ICSI), which was initially used to address severe male infertility, has replaced IVF as the most commonly used method for ART-mediated fertilization in many countries^3^. This more invasive fertilization procedure introduces additional mechanical damage, bypasses the complicated process of sperm-egg recognition and alters a series of downstream reactions^4^. Embryo cryopreservation enables embryos to be preserved for further transplantation, but both cryogens and freeze-thawing operation may cause damage to embryos^5^. All those unfavorable factors have raised concerns regarding the long-term health of ART-conceived children in recent years^6^. Despite claims to the contrary^7-9^, accumulating evidences have linked ART with potentially increased risks of neurodevelopmental disorders, cardiovascular dysfunction and metabolic abnormity in offspring and preeclampsia during pregnancy^10,11^.

Epigenetic modifications, such as DNA methylation and histone modifications, play key roles in regulating gene expression and are relatively sensitive to environmental factors^12^. Preimplantation embryos undergo dramatic genome-wide epigenetic reprogramming^13,14^ that coincides with the time frame of ART treatments. Thus, ART-associated perturbations may disturb the establishment and maintenance of epigenomic patterns and increase the relevant health risks of ART-conceived children in later life^15^. Notwithstanding, published researches on the association between ART and DNA methylation in offspring are limited to either specific genes^16,17^ or repeated sequences^18^ and restricted by methods^19-23^. Meanwhile, few studies have parsed the parental inheritance bias where considerable reports have revealed that parental genetic backgrounds, health situations, nutritional conditions and living habits have potential impacts on the epigenomes of neonates^24,25^. In addition, genome-wide changes in histone modifications of ART-conceived infants have not been reported and the effects of each specific ART procedure have not been fully elucidated so far.

Here, we integrated genome-wide maps of DNA methylation, four histone modifications associated with promoter/enhancer function (H3K4me1, H3K4me3, H3K27ac and H3K27me3), and gene expression for nuclear families to investigate the specific multilayer effects of various ART procedures on epigenomes in offspring. We found that various ART treatment would not dramatically disturb the global epigenome of neonates but subtly induced local and functional changes. Our comprehensive analysis not only accords with the findings in previous epidemiology studies but also reveals unexplored healthy risks in offspring from epigenomic aspect, and may serve as a valuable resource for researchers on the epigenetic influences of ART procedures.

## Results

### Global epigenomic profiles in ART-conceived neonates

To systematically study the DNA methylomic effects of different types of ARTs on offspring, we performed reduced-representation bisulfite sequencing (RRBS) on 137 umbilical cord blood (UCB) samples and 158 parental peripheral blood (PPB) samples from nuclear families with either singletons or twins (Fig. 1a, Supplementary Figure. 1a and Supplementary Table 1). The samples were classified into five groups based on the mode of conception: spontaneous (CTRL), IVF-ET, IVF-FET, ICSI-ET and ICSI-FET. There were no significant differences in the clinical features of the neonates among the different groups; in addition, maternal ages were under 35 years old and parental body mass index (BMI) values were comparable in general (Supplementary Table 2). In addition, we performed chromatin immunoprecipitation sequencing (ChIP-seq) on 33 UCB samples to examine histone modifications (H3K4me1, H3K4me3, H3K27ac, and H3K27me3) and mRNA sequencing (mRNA-seq) on 32 UCB samples to examine the transcriptomes of neonates (Fig. 1a; Supplementary Figure. 1a and Supplementary Table 1).

**Fig. 1.**
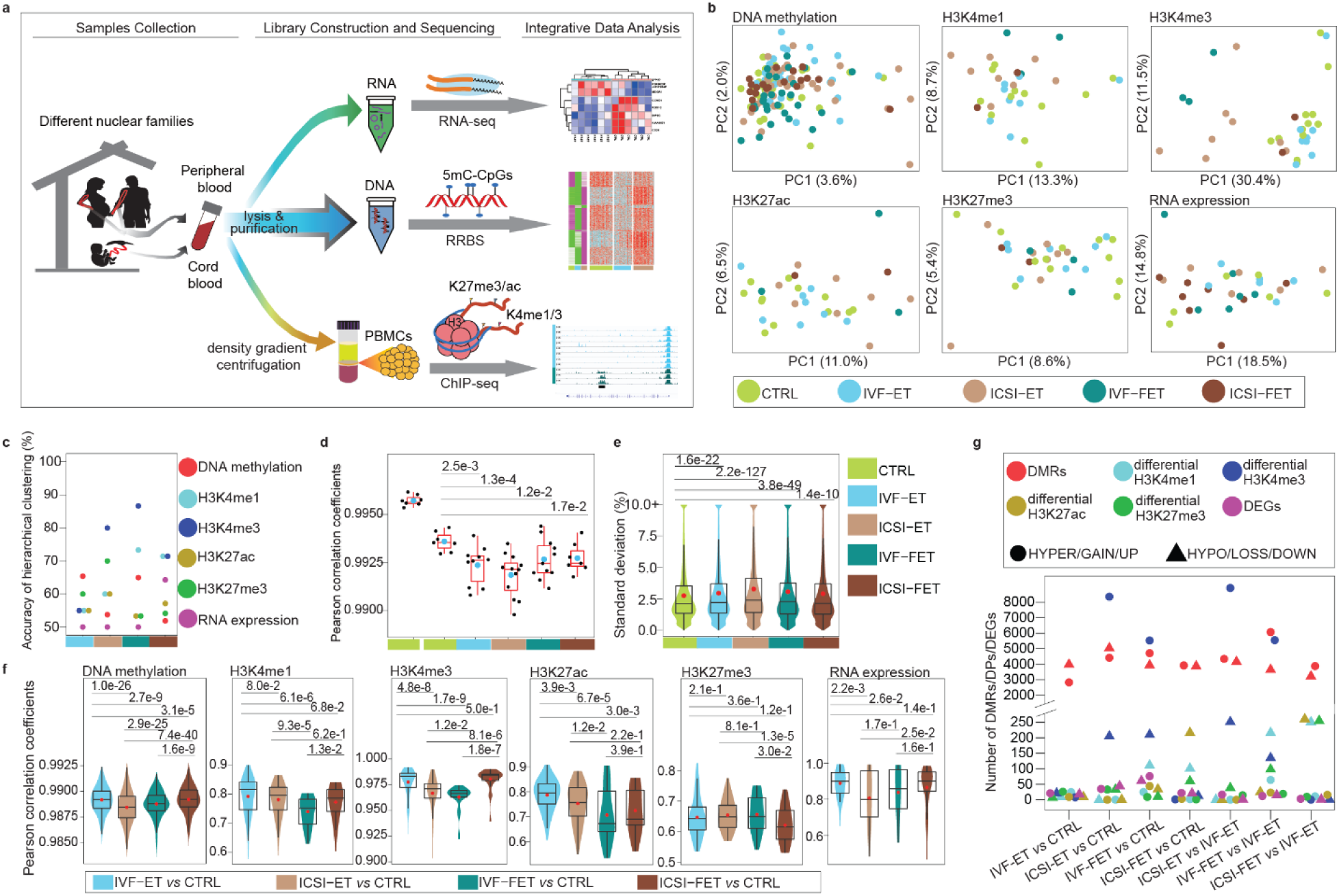
Epigenome profiling in ART-conceived neonates. **a.** Graphical overview of the study design**. b.** Principal component analysis (PCA) for all the five groups (CTRL, IVF-ET, ICSI-ET, IVF-FET, and ICSI-FET) of neonatal samples by using each layer of the reference epigenome and transcriptome. For DNA methylation, only the 100-bp tiles covered all neonatal samples were used; ChIP-seq peaks of all neonatal samples for each histone modification were pooled together, then the overlapping peaks were merged; genes with TPM=0 in all neonatal samples are removed. Finally, the number of genomic regions for each layer to generate the PCAs: DNA methylation (467097 100-bp tiles), H3K4me1 (195570 peaks), H3K4me3 (86157 peaks), H3K27ac (324900 peaks), H3K27me3 (257175 peaks), and RNA expression (26326 genes). **c.** The accuracy of hierarchical clustering for IVF-ET vs CTRL (cyan), ICSI-ET vs CTRL (sienna), IVF-FET vs CTRL (dark cyan), and ICSI-FET vs CTRL (dark sienna) neonatal samples. Samples and genomic regions used were the same as in **b. d.** Box plots for the distribution of the within-twin-pair Pearson correlation coefficients for genomewide DNA methylation in each group as in **b**, respectively. Twin pairs in CTRL group were classified into two subgroups, monozygotic and dizygotic twins. Each black dot represents the Pearson correlation coefficient and the cyan dots are the arithmetic means. The p-value between two groups was determined by unpaired and two-tailed t-test. **e.** Violin-box plots showed the distribution of standard deviations of genomewide DNA methylation for neonatal samples in each group mentioned in **b**, respectively. The p-value between two groups was determined by Wilcoxon rank-sum test. **f.** For each layer of the reference epigenome and transcriptome, violin-box plots showing the distribution of Pearson correlation coefficients between CTRL and one of the four ART groups. The red dots are the arithmetic means. The p-value between two groups was determined by Wilcoxon rank-sum test. **g.** The number of DMRs, DPs of four histone modifications, and DEGs for neonatal samples were shown for seven comparisons: IVF-ET versus CTRL, ICSI-ET versus CTRL, IVF-FET versus CTRL, ICSI-FET versus CTRL, ICSI-ET versus IVF-ET, IVF-FET versus IVF-ET, and ICSI-FET versus IVF-ET.

The results showed that the global DNA methylation levels, histone modifications, and transcriptomes of individual neonates were overall similar among the CTRL and four ART groups (Supplementary Figure. 1b-d). There were also no noticeable differences in global DNA methylation on various functional genomic regions, such as promoters, enhancers and repeats (Supplementary Figure. 1e). Unsupervised clustering of each layer of the reference epigenome showed no obvious subgroups but rather showed broadly distributed patterns among neonates, except for H3K4me3 in the IVF-FET and ICSI-ET groups (Fig. 1b and Supplementary Figure. 2a). These observations were further verified in hierarchical clustering analyses of the CTRL group versus specific ART subtype groups or ARTs as a whole; the accuracies in these analyses were close to a completely random value (50%) for binary classification but were markedly higher with regard to H3K4me3 in the IVF-FET and ICSI-ET groups (Fig. 1c and Supplementary Figure. 2b-h). These results suggested that ART processes do not dramatically disturb the overall epigenomes and transcriptomes of neonates in general but that H3K4me3 might be exceptionally sensitive to disturbance by specific ART procedures.

Correlation analysis for the DNA methylomes of twin pairs showed an overall high correlation coefficient greater than 0.99 (Fig. 1d); the coefficient for monozygotic twins was significantly higher than that for dizygotic twins in the CTRL group, consistent with previous studies^26,27^. The relatively lower correlation coefficients in all four ART subgroups than in the CTRL group implied that ART processes might subtly affect the epigenomes of neonates (Fig. 1d). Additionally, this phenomenon could also be observed when only same-sex twins were analyzed or in any two non-twin-pair neonates in each group (with exclusion of twin-pair bias) (Supplementary Figure. 3a-c). Moreover, the standard deviations of the DNA methylation levels of all neonates in the different ART groups were slightly but significantly higher than that in the CTRL group (Fig. 1e). Above results indicated that ART itself might increase heterogeneity in neonates. Intergroup correlation analysis revealed that the IVF-ET group was the most similar to the CTRL group with regard to H3K4me1, H3K4me3, H3K27ac, transcriptomes and DNA methylomes genome-wide or on special elements (Fig. 1f and Supplementary Figure. 3d), indicating that IVF-ET might have less epigenetic influence on neonates than the other three ART processes. These findings suggested that imperceptible disturbances might be introduced into neonates and might vary among different ART subtypes.

### Subtle epigenomic changes in ART-conceived offspring

To elucidate the epigenetic effects of ARTs on neonates, we performed pairwise comparisons between groups for DNA methylation, histone modifications, and gene expression. In particular, to identify differentially methylated regions (DMRs), six comparisons were generated using all neonates, only singleton neonates, and four groups of neonates from twin cohorts. DMRs in at least four comparisons were selected for downstream analysis to improve the confidence of the analysis (Supplementary Figure. 3e, see Methods for more details). The numbers of hyper-/hypo-DMRs, gain/loss differential histone modifications (differential peaks, DPs), and up-/downregulated differentially expressed genes (DEGs) among the different comparison groups are shown in Fig. 1g and Supplementary Table 4 (see Supplementary Table 5-10 for more details). The difference detected in DMRs was mainly less than 15%, which was relatively moderate (Supplementary Figure. 4a). Meanwhile, only a few DPs were observed in the comparison between the IVF-ET and CTRL groups for four histone modifications, suggesting that IVF-ET has little impact on histone modification (Fig. 1g and Supplementary Table 4). The numbers of H3K4me3 DPs detected in the comparison between the ICSI-ET/IVF-FET and CTRL groups (ICSI-ET: 8352 gain, 205 loss; IVF-FET: 5526 gain, 210 loss) were consistent with the observation in unsupervised clustering (Fig. 1b and Supplementary Figure. 2a, f-g), while the majority of the increased DPs in the ICSI-ET/IVF-FET groups were in regions with originally weak signals in the CTRL and IVF-ET groups (Supplementary Figure. 4b). All of the DEGs, DMRs and DPs for each ART group versus the CTRL group were further validated at the individual level and broadly distributed on the genome scale (Supplementary Figure. 4c-e; Supplementary Figure. 5). Together, these results indicated that ART processes potentially caused subtle epigenetic changes distributed widely throughout the genome.

### Parent-derived and ART-derived DNA methylomic differences in offspring

To evaluate parental influence on the DNA methylomes of progeny, the DNA methylation levels of DMRs identified in neonates were further analyzed in parents. There were significant differences between the two groups of fathers or mothers (Fig. 2a; Supplementary Figure. 6a; p<0.01). In addition, approximately 50% of DMRs identified in neonates significantly overlapped with DMRs identified in their corresponding parents (Fig. 2b and Supplementary Figure. 6b, Supplementary Table 14). To investigate the effects of ARTs *per se*, the overlapping DMRs were removed and the remaining neonatal DMRs were used as the final neonatal DMRs for downstream analysis (Supplementary Figure. 6c, Supplementary Table 11 and 15). As expected, no significant differences were observed in the DNA methylation levels of the final neonatal DMRs between the corresponding parental groups (Supplementary Figure. 6d-e).

**Fig. 2.**
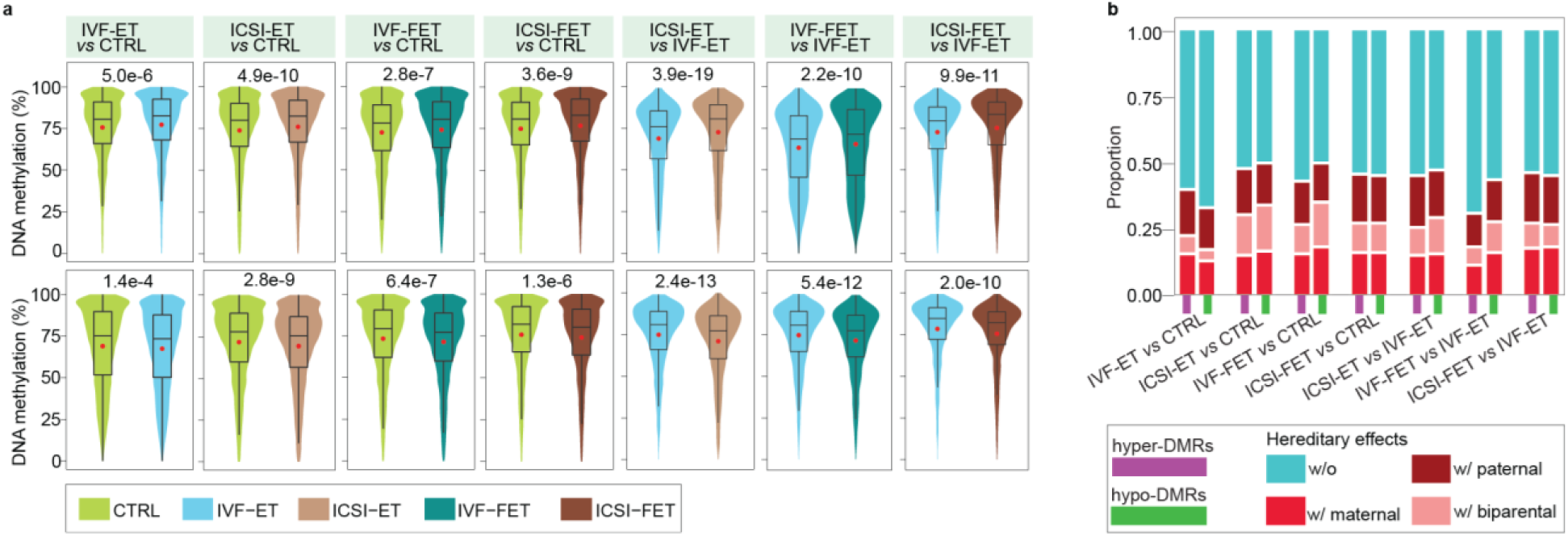
Remove the parental effects in neonatal DMRs. **a.** Violin-box plots displayed the distribution of DNA methylation level in paternal samples at hyper-(upper) and hypo-DMRs (lower) of the seven comparisons, IVF-ET versus CTRL, ICSI-ET versus CTRL, IVF-FET versus CTRL, ICSI-FET versus CTRL, ICSI-ET versus IVF-ET, IVF-FET versus IVF-ET, and ICSI-FET versus IVF-ET neonatal samples. The p-value between two groups was determined by Wilcoxon rank-sum test. **b.** Bar graph showed the proportion of DMRs with/without hereditary effects in the hyper- or hypo-DMRs of each comparison. Any one of the 14 groups of DMRs were classified into four subgroups, without hereditary effects (w/o), only with paternal hereditary effects (w/paternal, only overlapped with the DMRs of comparison for the corresponding fathers), only with maternal hereditary effects (w/ maternal, only overlapped with the DMRs of comparison for the corresponding mothers), and with biparental hereditary effects (w/ biparental, overlapped with the DMRs of comparisons for the corresponding fathers and mothers).

### Epigenomic alteration of regulatory regions in IVF-ET-conceived offspring

IVF-ET, the most basic process of ART treatment, is generally recommended as a first-line ART therapy for couples with female infertility^28^. A total of 1703 hyper-DMRs and 2658 hypo-DMRs were identified in the IVF-ET group compared with the CTRL group (Supplementary Table 11). Gene Ontology (GO) enrichment analysis showed that the associated genes of DMRs were enriched in a broad range of processes, including processes related to the nervous system, respiratory system and cardiovascular system, etc. (Fig. 3a; Supplementary Table 16). Human Phenotype Ontology (HPO) analysis also indicated that alterations in DNA methylome induced by IVF-ET might lead to abnormal phenotypes in ocular, cardiovascular and skeletal system, tooth morphology and metabolism, etc. (Fig. 3b; Supplementary Table 16).

**Fig. 3.**
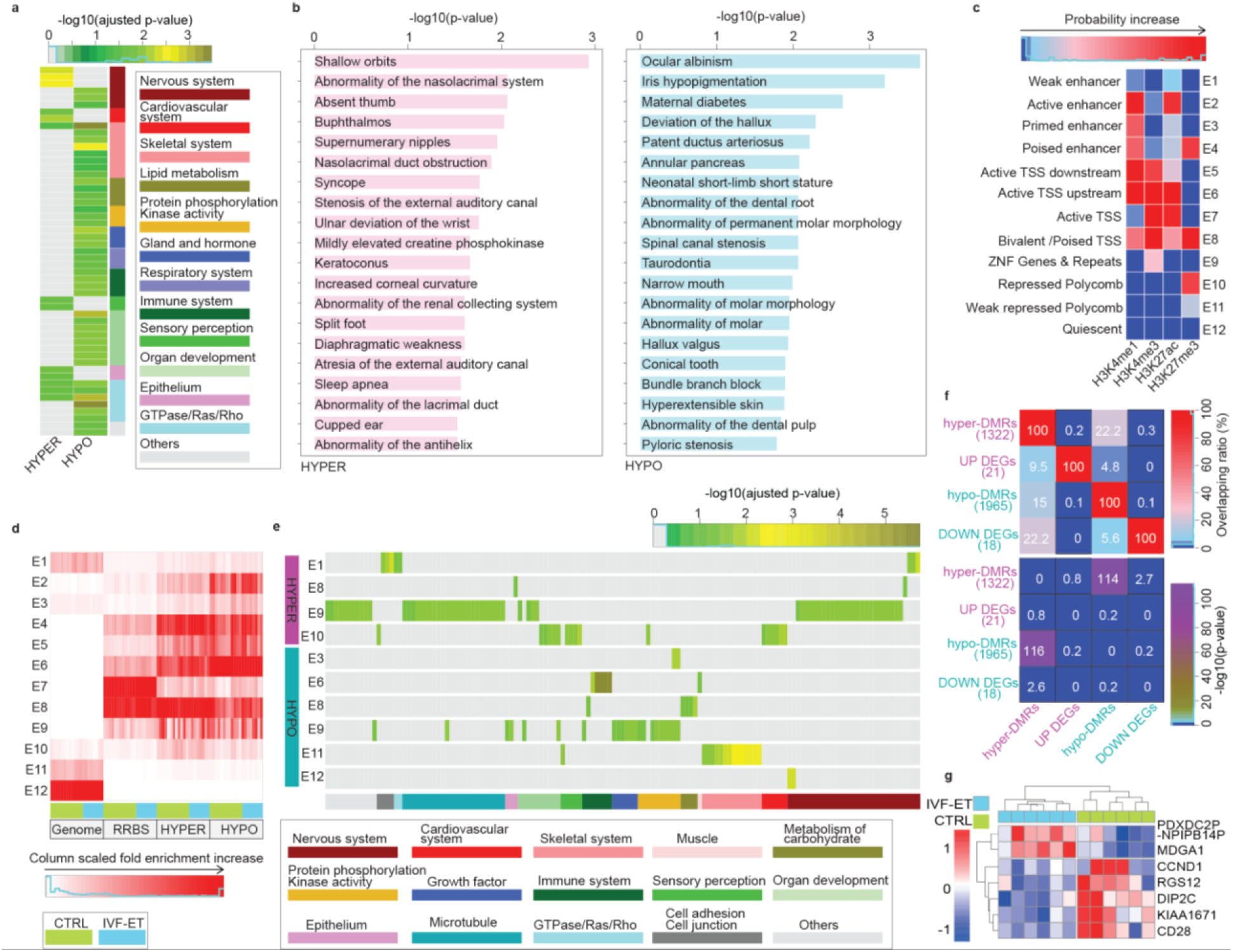
IVF effects on the epigenome of the offspring. **a.** Gene ontology (biological process) analysis for the hyper- and hypo-DMRs of IVF-ET versus CTRL, all enrichment terms with adjusted p-value < 0.05 were grouped and shown. **b.** The top 20 enriched ontology terms of human phenotype for hyper-(left) and hypo-DMRs (right). **c.** Based on the four histone modifications in all neonatal samples, 12 chromatin states were defined by chromHMM. Each row of the heatmap corresponds to a specific chromatin state, and each column corresponds to a different histone mark, H3K4me1, H3K4me3, H3K27me3, or H3K27ac. **d.** In each neonatal sample of CTRL and IVF-ET groups, column scaled fold enrichment of the chromatin states at genomewide regions (Genome), RRBS covered regions (RRBS), hyper- and hypo-DMRs (HYPER and HYPO) was shown. **e.** For each subgroup of DMRs in **d**, its gene ontology (biological process) enrichment analysis results were grouped and shown, if its enrichment terms existed (adjusted p-value < 0.05). **f.** Heatmap showed overlapping ratio among associated genes of DMRs and DEGs for IVF-ET vs CTRL (upper), and their statistical significances (lower) determined by hypergeometric test. **g.** Heatmap showed the expression level of DEGs overlapped with the associated genes of DMRs (PDXDC2P-NPIPB14P, MDGA1, CCND1, RGS12, DIP2C, KIAA1671, CD28) among neonatal samples in CTRL and IVF-ET.

Given the distinct roles of various genomic regions in regulating gene expression, we performed ChromHMM using four types of histone modifications and identify 12 chromatin states to investigate the specific impacts of IVF-ET on the DNA methylation of functional elements (Fig. 3c). The regions covered in our DNA methylation data were mainly the regions related with transcription start site (TSS) (E7 and E8), consistent with the features of highly enriched CpG regions in RRBS. Notably, a large fraction of DMRs were concentrated on Active enhancer (E2), Poised enhancers (E4), Active TSS upstream sites (E6) and Bivalent/Poised TSSs (E8), but almost depleted from Active TSSs (E7) (Fig. 3d). GO analysis for those DMRs in different chromatin states suggested that those hyper-DMRs in Weak enhancer (E1), Bivalent/Poised TSSs (E8) and ZNF Genes & Repeats (E9) might be associated with the interference on nervous system, while hyper-DMRs in Repressed Polycomb (E10) might account for the influence on cardiovascular system. Meanwhile, hypo-DMRs in Active TSS upstream (E6) and Weak repressed polycomb (E11) might highly correlated with immune system and skeletal system, respectively (Fig. 3e, Supplementary Table 16). In fact, the intersection of the DMR associated genes with the DEGs revealed that the majority of the DMRs were not associated with transcriptional changes in their associated genes (Fig. 3f, Supplementary Table 17). For DEGs overlapped with hypo-DMRs associated genes, *MDGA1* was upregulated in the IVF-ET group, while *RGS12* was downregulated; in the former, the DMRs occurred in promoters, while in the latter they occurred in introns. The expression of genes associated with hyper-DMRs in introns or distal intergenic regions, such as *KIAA1671* (introns), *DIP2C, CCND1* and *CD28*, was decreased (Fig. 3g). Polymorphisms/mutations in *MDGA1* and *DIP2C* are associated with mental disease, and dysregulation of *KIAA1671, CCND1*, and *DIP2C* has been reported to take part in tumorigenesis; furthermore, *RGS12* dysfunction contributes to tumorigenesis as well as pathological cardiac hypertrophy, osteoclast genesis and bone destruction. Together, our findings indicated that DNA methylomic changes caused by IVF-ET might affect multiple aspects related with neonates’ health, but mainly kept away from active TSS regions.

### Identification of altered epigenomic profiles associated with ICSI procedures

To investigate the differences between those two ways of fertilization, we first compared ICSI-ET with IVF-ET and identified more than 4500 DMRs and 9000 H3K4me3 DPs (Supplementary Tables 4 and 11; 2365 hyper-DMRs and 2386 hypo-DMRs; 8914 gain DPs and 270 loss DPs). Although there were no overlapped genomic locations, the changes shared common associated genes and GO terms (Extended Data Figs. 7a-c), with only a few genes showed consistent changes in expression with changes in the epigenome (Supplementary Tables 18). Taking the differences between the ICSI-ET and CTRL groups into account enabled us to determine the abnormal effects of ICSI *per se*. Thereafter, the DMRs and H3K4me3 DPs of the ICSI-ET group versus the IVF-ET group were refined, the former based on k-means clustering of DNA methylation levels among the ICSI-ET, IVF-ET and CTRL groups and the latter based on integrated comparison between DPs of ICSI-ET vs CTRL (n=8557) and DPs of ICSI-ET vs IVF-ET (n=9184). Six clusters of DMRs (C1∼C3 for hyper-DMRs, C4∼C6 for hypo-DMRs) and overlapping H3K4me3 DPs (n=7317) that showed similar change tendencies in the comparison of ICSI-ET vs IVF-ET and ICSI-ET vs CTRL groups were selected for downstream analysis (Fig. 4a-b; Extended Data Figs. 7d; Supplementary Tables 19). The associated genes of hyper-DMRs and gain DPs were enriched in processes involving the development of skeletal system and Wnt signaling morphogenesis, respectively (Fig. 4c; Supplementary Table 20). Meanwhile, HPO annotation of those DMRs also revealed that the disturbance in those regions might potentially introduce adverse effect on appendage and skeletal system (Extended Data Figs. 7e; Supplementary Table 20). Though there were only 84 loss DPs of H3K4me3, they were highly enriched in GO terms of immune system (Fig. 4c; Supplementary Table 20). Similar to the observation in IVF-ET, the selected DMRs for ICSI also tended to avoid the active TSSs (Fig. 4d). Associated genes of hyper DMRs in ZNF Genes & Repeats (E9) and weak repressed polycomb (E11) were enriched in the processes of immune and nervous system, respectively. Associated genes of hypo-DMRs in Active TSS (E7) and ZNF Genes & Repeats (E9) were also enriched in the processes of immune and nervous system, respectively (Fig. 4e; Supplementary Table 20). As expected, H3K4me3 peaks were mainly enriched in regions around TSSs (E5, E6, E7, E8). However, regions for the selected gain H3K4me3 DPs in CTRL and IVE-ET group were highly enriched in active enhancer (E2) and poised enhancer (E4) and shifted to ZNF Genes & Repeats (E9) in ICSI-ET. For loss H3K4me3 DPs, the situation was reversed (Fig. 4d). GO terms for the selected gain H3K4me3 DPs in E2 and E4 were mainly related to the regulation of leukocytes, cell migration/adhesiveness and the development of several systems (Fig. 4e; Supplementary Table 20). These results suggested that ICSI might alter H3K4me3 at enhancers and affect multiple processes. Together, the results regarding both the DNA methylome and H3K4me3 modification raise the question of whether ICSI may have a potential impact on health of offspring, with the skeletal systems and immune system as the representatives.

**Fig. 4.**
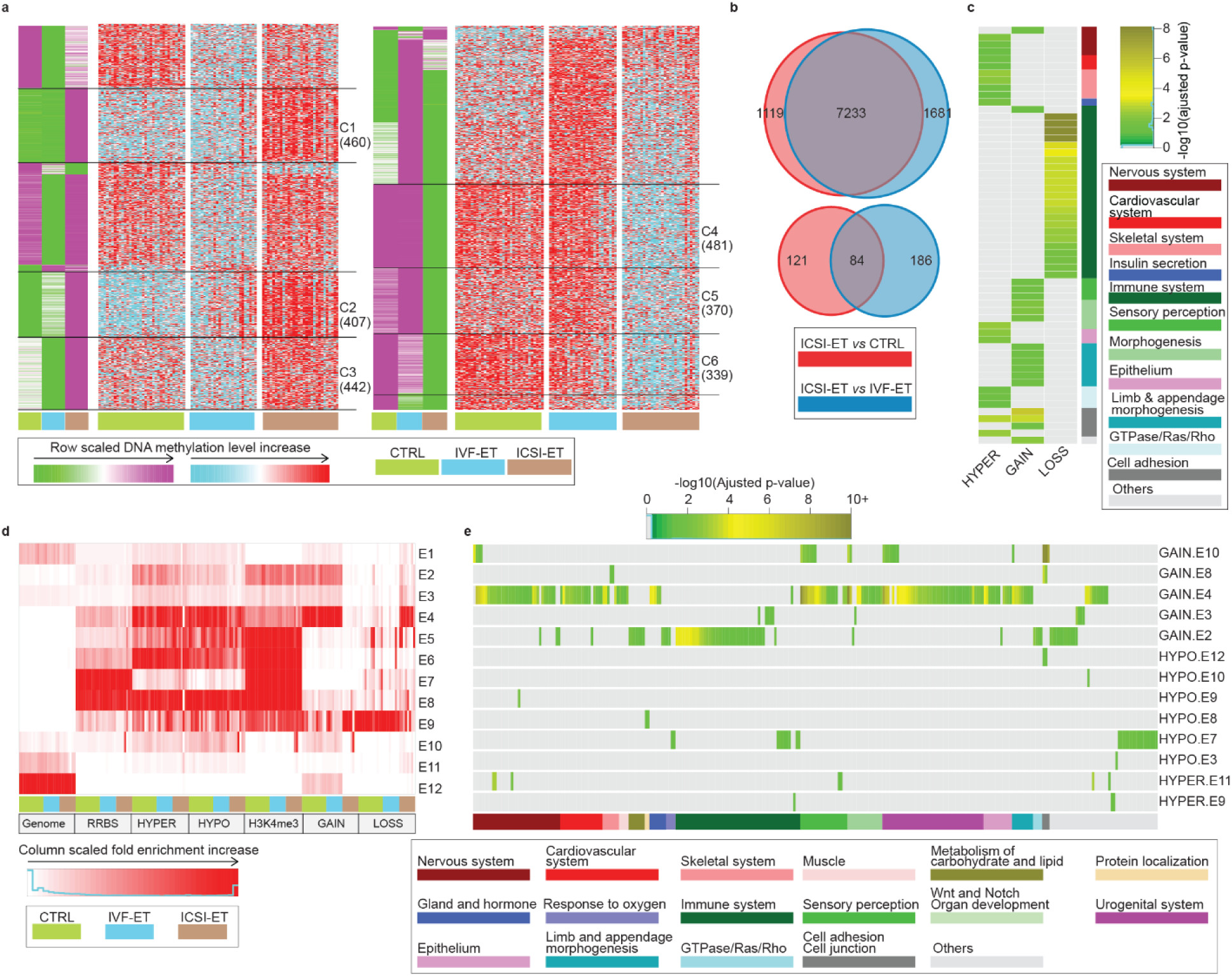
Separate epigenetic effects of ICSI. **a.** K-means clustering for the hyper-(left) and hypo-DMRs (right) between ICSI-ET vs IVF-ET neonatal samples. In green-purple heatmaps, the hyper-DMRs (hypo-DMRs) were classified into seven (nine) clusters by k-means clustering on row scaled average DNA methylation level. With the same row order, row scaled DNA methylation level in each neonatal sample was also shown in cyan-red heatmap. The number of DMRs in the selected clusters were also shown. **b.** Venn diagram for the overlap between of the gain (upper) and lost (lower) H3K4me3 DPs of ICSI-ET vs CTRL, and ICSI-ET vs IVF-ET neonatal samples. **c.** For the selected hyper-(union of cluster C1-C3) and hypo-DMRs (union of cluster C4-C6), gain (7233) and loss (84) H3K4me3 DPs, their gene ontology (biological process) enrichment analysis results were grouped and shown, if its enrichment terms existed (adjusted p-value < 0.05). **d.** The chromatin states distribution of the selected hyper- and hypo-DMRs and H3K4me3 DPs (7233 GAIN, 84 LOSS) in each neonatal sample of CTRL, IVF-ET, and ICSI-ET. The chromatin states were defined in **fig. 3c. e.** For DMRs or DPs in different states in **d**, their gene ontology (biological process) enrichment analysis results were grouped and shown, if the enrichment terms existed (adjusted p-value < 0.05).

### Specific epigenomic changes induced by freeze-thawing procedure

We then investigated the differences between the IVF-FET and IVF-ET groups and observed 6251 DMRs and 5683 DPs of H3K4me3 (Supplementary Tables 4 and 11; 4191 hyper-DMRs and 2060 hypo-DMRs; 5548 gain DPs and 135 loss DPs). In addition, compared to the number of DPs in the ICSI-ET group versus the IVF-ET group or the CTRL group, the notably increased numbers of H3K4me1, H3K27me3 and H3K27ac DPs suggested that the freeze-thawing procedure might introduce more disturbance of histone modifications than ICSI (Supplementary Figure. 8a; Supplementary Table4). Moreover, the DMRs and DPs (of H3K4me3 and H3K27ac) shared common associated genes and GO terms, implying that freeze-thawing procedure might cause certain functional changes through different epigenetic layers (Fig. 5a and Supplementary Figure. 8b; Supplementary Table 21∼22).

**Fig. 5.**
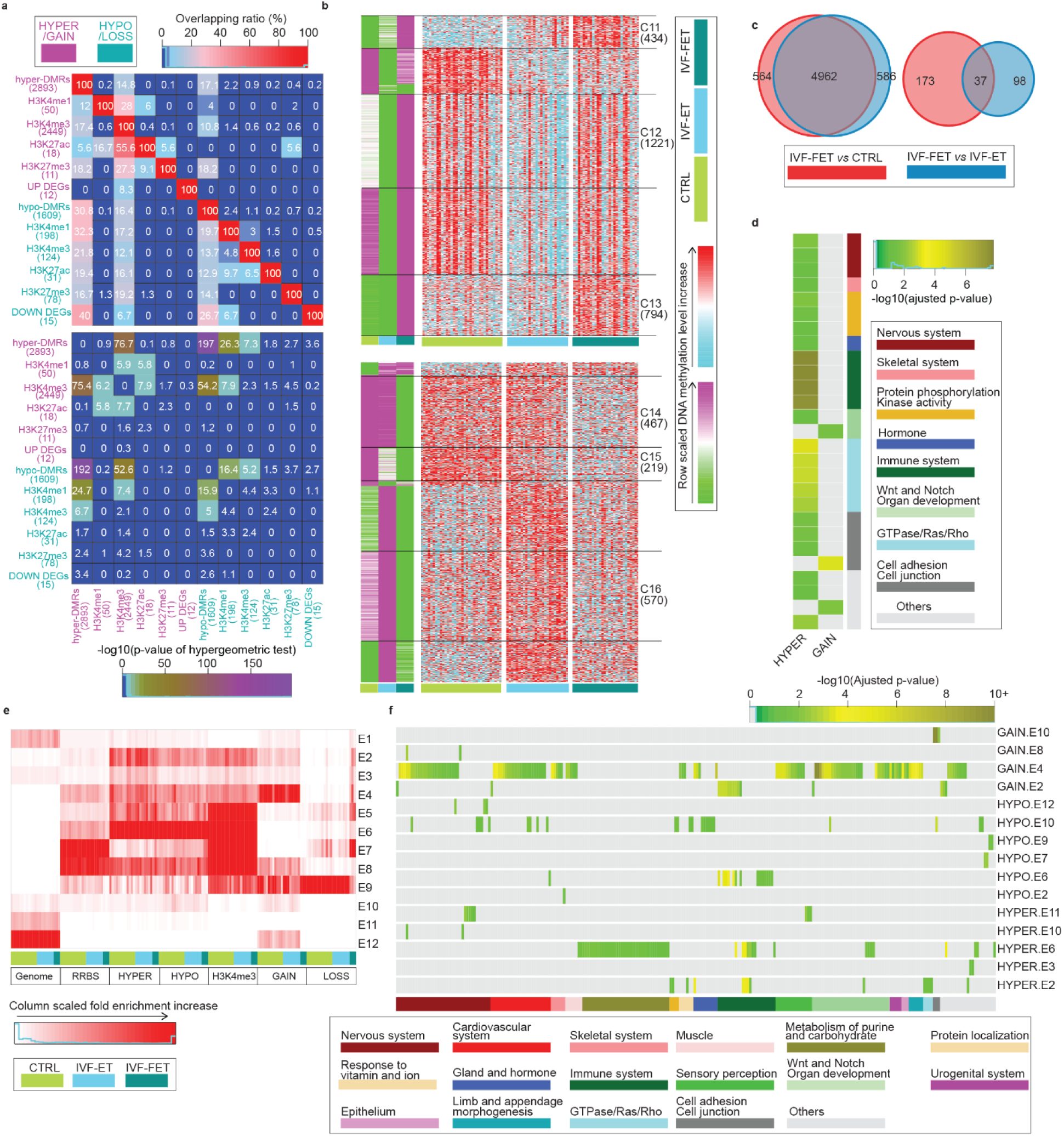
Separate epigenetic effects of freezing-thaw. **a.** Overlapping ratio among DEGs, associated genes of DMRs and DPs for IVF-FET vs IVF-ET neonatal samples (upper), and their statistical significances (lower) determined by hypergeometric test. **b.** K-means clustering for the hyper-(upper) and hypo-DMRs (lower) between IVF-FET vs IVF-ET neonatal samples. In green-purple heatmaps, the hyper-DMRs (hypo-DMRs) were classified into six (seven) clusters by k-means clustering on row scaled average DNA methylation level. With the same row order, row scaled DNA methylation level in each neonatal sample was also shown in cyan-red heatmap. The number of DMRs in the selected clusters were also shown. **c.** Venn diagrams displayed the overlap between the gain (left) and lost (right) H3K4me3 DPs of IVF-FET vs CTRL and that of IVF-FET vs IVF-ET. **d.** For the selected hyper-(union of cluster C11-C13) and hypo-DMRs (union of cluster C14-C16), overlapping gain (4962) and loss (37) H3K4me3 DPs, their gene ontology (biological process) enrichment analysis results were grouped and shown, if the enrichment terms existed (adjusted p-value < 0.05). **e.** The chromatin states distribution of the selected hyper- and hypo-DMRs, overlapping 4962 gain and 37 loss H3K4me3 DPs in each neonatal sample of CTRL, IVF-ET, and IVF-FET. The chromatin states were defined in **fig. 3c. f.** For DMRs or DPs in different states in **e**, all enrichment terms of gene ontology (biological process) with adjusted p-value < 0.05 were grouped and shown.

To further explore the specific effects of freeze-thawing procedure *per se*, we selected six subgroups of DMRs of IVF-FET vs IVF-ET through k-means clustering of scaled DNA methylation levels among the IVF-FET, IVF-ET and CTRL groups (Fig. 5b; Supplementary Table 23; cluster C11∼13 in hyper-DMRs and cluster C14∼C16 in hypo-DMRs), as well as H3K4me3 DPs (n=4999) identified by intersection in the comparison of IVF-FET vs IVF-ET (n=5683) and IVF-FET vs CTRL (n=5736) (Fig. 5c). The replications of DMRs and DPs for each group were highly comparable (Supplementary Figure. 8c). The associated genes of selected hyper-DMRs were highly enriched in the processes of neutrophil-mediated immunity and the regulation of GTPase/Ras (Fig. 5d). The results of HPO analysis further revealed that the selected DMRs might be related to carcinogenesis (Supplementary Figure. 8d; Supplementary Table 22). The majority of selected hyper/hypo-DMRs were inclined to evade from Active TSS (Fig. 5e), and the associated genes of those DMRs in Active enhancer (E2) and Active TSS upstream (E6) were highly enriched in the processes of immune system, metabolism of purine/carbohydrate, muscle and skeletal system (Fig. 5f; Supplementary Table 22). Poised enhancer (E4) was the most overrepresented states for the selected gain H3K4me3 DPs in CTRL/IVF-ET group and became ZNF Genes & Repeats (E9) in IVF-FET. Associated genes for the selected gain H3K4me3 DPs in E2 and E4 were enriched in the processes of certain systems (Fig. 5e-f; Supplementary Table 22). It was also worth mentioning that the genes *PGP, PLOD3*, and *STAB1* in the IVF-FET group all exhibited decreased expression with hyper-DMRs in their promoters. In addition, the expression level of *SEPT9* also showed decreased tendency in the IVF-FET group, with hyper-DMRs detected in the TSS upstream region and hypo-DMRs detected in the TSS downstream region (Supplementary Table 21). Considering that the dysregulation of the *PGP, PLOD3, STAB1* and *SEPT9* genes as well as the GTPase signaling pathway is potentially associated with carcinogenesis^29^, the above results implied that disturbance in the epigenome introduced by cryopreservation *per se* might potentially affect the immune system and increase the risk of cancer later in life.

### Common epigenetic effects among different ART processes

Since various ART processes share common procedures, we further analyzed the relationship among DMRs/DPs to investigate the common epigenetic effects of ART treatments. Clearly, the DMRs identified in seven pairwise comparisons were overlapped significantly in genomic location and shared many common GO terms (Supplementary Figure. 9a-b; Supplementary Table 24). Moreover, a considerable number of DMRs showed similar change tendencies in different ART groups compared with CTRL group through k-means clustering using DMRs from all seven groups (Fig. 6a; Supplementary Table 25; C22, n=931; C23, n=776). The associated genes of C22 and C23 were highly enriched in the regulation of GTPase activity, stress-activated MAPK cascade, cell morphogenesis and neuron projection development in GO analysis (Fig. 6b; Supplementary Table 24).

**Fig. 6.**
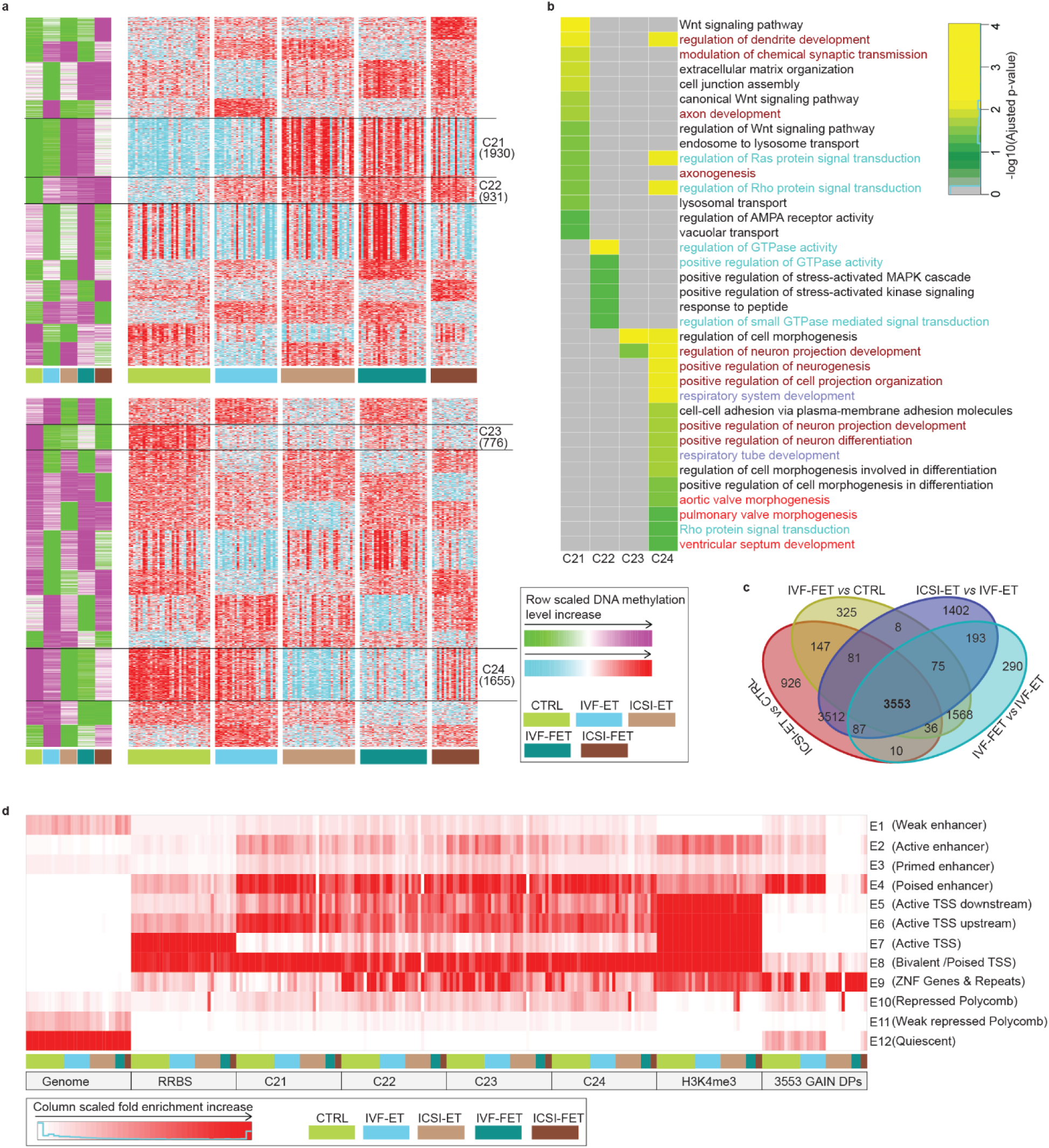
Common Epigenetic effect of different ARTs. **a.** K-means clustering for the all merged hyper-(up) and hypo-DMRs (down) of the seven comparisons. The hyper- and hypo-DMRs of the seven comparisons were merged if they have 1-bp common region at least, respectively. In green-purple heatmaps, DMRs were classified into twelve clusters by k-means clustering on row scaled average DNA methylation level. With the same row order, row scaled DNA methylation level in each neonatal sample was also shown in cyan-red heatmap. The number of DMRs in the selected clusters were also shown. **b.** For the selected clusters of hyper-DMRs (C21, C22) and hypo-DMRs (C23, C24), all enrichment terms of gene ontology (biological process) with adjusted p-value < 0.05 were grouped and shown. **c.** Venn diagrams displayed the overlap of the gain H3K4me3 DPs for ICSI-ET vs CTRL, IVF-FET vs CTRL, ICSI-ET vs CTRL, and IVF-FET vs IVF-ET. **d.** The chromatin states distribution in each neonatal sample for the selected DMRs and overlapping gain H3K4me3 DPs. The chromatin states were defined in **fig. 3c**.

Interestingly, clusters C21 and C24 seemed to represent regions where ICSI and freeze-thawing procedure might introduce similar alterations, revealed by the similar change on DNA methylation levels in the ICSI-ET and IVF-FET groups compared with CTRL and IVF-ET groups (Fig. 6a, Supplementary Table 25). These DMRs were related to Wnt signaling pathway, respiratory, cardiovascular and nervous system. Similarly, there were noticeable overlaps between ICSI- and freeze-thawing-specific gain H3K4me3 DPs (Fig. 6c; Supplementary Table 25; C1-C3 and C11-C13; n=3553), which were mainly involved in immune (B lymphocytopenia, Abnormality of B cell number, Neutropenia) and skeletal system by HPO analysis (Supplementary Figure. 9c; Supplementary Table 24). As mentioned previously, those DMRs were also noticeably absent from Active TSS (E7), with the increased enrichment in ZNF Genes & Repeats (E9) for DMRs in clusters C21 and C24 (Fig. 6d). Meanwhile, the difference between ICSI-FET and IVF-ET would be expected as a mixed effect of ICSI and freeze-thawing operation, indeed, large percentage of those DMRs showed similar changes in ICSI-ET or IVF-FET (C31∼C33; C35, C36, C38). However, 13% hyper-DMRs (C34) and 19% hypo-DMRs (C37) were uniquely found in ICSI-FET groups, implying complex interactions between ICSI and freeze-thawing (Supplementary Figure. 9d, Supplementary Table 25). The above results together suggested that in addition to causing their own unique effects, as mentioned previously, distinct ART procedures also leaded to certain common effects on the epigenomes of offspring.

## Discussion

Safety concerns regarding ARTs are as old as ARTs themselves^30^. Although a number of studies suggest that ART may have adverse effects on the long-term health on offspring, the underlying mechanisms remain to be elucidated. We systematically explored the influences of fertilization procedures and freeze-thawing operation on descendant genome-wide DNA methylation, histone modifications and gene expression. Recruitment of nuclear families with twins provided us with perfect biological replicates in each family and enabled us to eliminate ART-irrelevant parental impacts as much as possible. Our study illustrates that the epigenomes of neonates conceived by ART are overall similar to those of naturally conceived children, which may partially explain why most ART offspring are generally healthy. However, ART do increase the heterogeneity of the DNA methylome within twin pairs and induce local subtle changes in different epigenetic layers, supporting the theory that ART increases risks of epigenomic abnormality^16-18^. We also found that more than half of the DNA methylomic changes in ART offspring were derived from parents, highlighting the necessity of removing parental bias in assessing the influences of ARTs *per se*. Moreover, we reported the genome-wide impacts of ARTs on four types of histone modifications in humans for the first time. The epigenomes of IVF-ET conceived infants seemed to be closer to those of naturally conceived infants than those of IVF-FET or ICSI-ET conceived infants and showed almost no disturbance in either type of histone modification, suggesting that ART-inherent procedures, including controlled ovarian hyperstimulation (COH), in vitro culture, in vitro fertilization, etc., may not increase the risks of abnormal histone reprogramming; however, ICSI and freeze-thawing operation may do so. What surprised us was that H3K4me3 was the most profoundly impacted by ICSI and freeze-thawing compared with the other three types of histone modifications (H3K4me1, H3K27me3 and H3K27ac), therefore, H3K4me3 might serve as a sensitive histone modification for assessment of the influences of ARTs.

The distributions of epigenetic changes tended to be away from active cis-regulatory elements and were not largely associated with transcriptional changes in corresponding genes. Functional enrichment of epigenetic changes suggested that epigenetic disorders caused by ICSI might interfere the processes associated with skeletal system, which is also highlighted in a recent study about the potential impact of ART on DNA methylome^31^. Our results also indicated that freeze-thawing procedure, as well as ICSI, might increase the risks of immune dysfunction in offspring. Though the risks of hospital admission in FET-conceived children and asthma medication in ART-conceived children have been reported to be higher than CTRL conceived children^32,33^, direct evidence regarding the effects of ART on the immune system is lacking so far and long-term follow-up studies are required. Previous studies have revealed elevated rates of preeclampsia in women who have undergone FET^34^. GTPases, especially Rho kinases, play essential roles in extravillous trophoblast cell (EVT) invasion^35^, and limited EVT invasion following poor remodeling of arteries is widely observed in preeclampsia^36^. More interestingly, our results suggested that freeze-thawing operation might cause dysregulation in the GTPase/Ras signaling pathway in offspring. Thus, the epigenetic abnormality we report may also possibly explain the increased risks of preeclampsia in FET compared with fresh ET pregnancies. Our findings emphasize that it is reasonable to inform patients of potential risks associated with ICSI and embryo cryopreservation and that it would be wise to reconsider overutilization of ICSI or routine freezing of all embryos.

Apart from the influences for given ART operations, we also revealed the common effects induced by various ART processes. Further analysis showed that ART-induced DNA methylation changes were enriched in the processes of cardiovascular system and glucolipid metabolism in the comparisons of all four ART groups versus CTRL groups. Since the abnormalities in lipid profiles and higher rates of cardiovascular dysfunction have been reported in ART-achieved children^11,37^, it implies that the disturbance on epigenome by ART in offspring may increase the risk of metabolic syndrome^38^. In addition, we identified a considerable number of common DMRs among different ART groups compared with CTRL group. It is noteworthy that the regulation of GTPase activity was the most overrepresented terms in GO analysis for those DMRs, which is in line with those studies suggesting that pregnancies conceiving by ART is related with the increased risk for certain cancers in offspring and preeclampsia compared with natural pregnancy^39,40^.The common DMRs induced by both ICSI and freeze-thawing procedure were enriched in the processes involving in neuron, consistent with the concern that these two kinds of aggressive ART operation might increase the risk of mental disorders in offspring^41^. As suggested by the reports that improved in vitro culture systems for animals will affect the epigenome less than earlier versions^42,43^, continuous optimization for ART procedures is urgently needed to simulate the in vivo environment and reduce potential epigenetic abnormalities in offspring.

Our study was mainly focused on the effects of different fertilization methods and freeze-thawing and only involved newborns after full-term pregnancy, lacking postnatal follow-up data on the enrolled populations. A larger multicenter randomized controlled trial (RCT) along with detection of the epigenetic profiles of offspring in later life would be helpful for elucidation of continued epigenetic change and exploration of the specific epigenetic impacts of other factors in ART treatment, such as the duration of embryo culture or the composition of the culture system.

In conclusion, our results provide an epigenetic basis for the increased long-term health risks in ART offspring. Our study highlights ART clinical interventions that require particular surveillance. More effort should be expended to optimize current ART systems, and the choice of appropriate procedures requires careful evaluation. Since epigenomic changes might be maintained throughout the human lifespan^44^ and can potentially be transmitted to subsequent generations, long-term follow-up and health evaluation of ART offspring are necessary to provide more robust clinical evidence.

## Methods

Methods and any associated references are available in the supplemental sections.

## Acknowledgments

We thank the families participated in this study. We thank Robert Norman for discussion and reviewing the manuscript. This project is funded by National Natural Science Foundation of China (81730038; 81521002), National Key Research and Development Program (2018YFC1004000; 2017YFA0103801; 2017YFA0105001) and Strategic Priority Research Program of the Chinese Academy of Sciences (XDA16020703). Y.W. was supported by Postdoctoral Fellowship of Peking-Tsinghua Center for Life Science.

## Author contributions

W.C., X.M., Y.P. and S.K. wrote the manuscript. W.C. and X.M. collected the study materials and samples and patient data, performed the experiments. Y.Wei., Y.Z., S.T. and W.Z. were helpful for the recruitment of family and sample collection for this study. Y.P. developed analysis methods and performed bioinformatic analysis. J.Q., L.Y. and Y.Wang. developed the experimental conception and designs. All the authors read and approved the final manuscript.

